# Most science is published from countries lacking in democracy and freedom of press

**DOI:** 10.1101/2025.07.03.663115

**Authors:** John P.A. Ioannidis, Jeroen Baas

## Abstract

**Background:** Democracy and freedom of press may affect how science is prioritized, produced, communicated and disseminated. We aimed to map the production of scientific publications worldwide in terms of democracy and freedom of press ratings of countries.

**Methods:** This is a bibliometric study cross-linking global bibliometric data with democracy ratings and freedom of the press indices for countries around the world. Democracy ratings used the Democracy Index in 2024 and in 2006 (when first released by the Economist Intelligence Unit) and Freedom of Press ratings used the 2024 index by Reports Without Borders. The Scopus database was used for publications from each country. Fractional counts were assigned for publications co-authored by authors from different countries. Full articles, reviews, conference papers, books and book chapters were included.

**Results:** In 2024, countries characterized as full democracies produced only 22% of the Scopus-indexed publications, versus 66% in 2006. There was no correlation between the ratio of publications indexed in 2024 versus 2006 and the absolute or relative change in Democracy Index between 2006 and 2024 (r=0.02 and r=0.00, respectively). 78% of publications in 2024 came from countries with problematic (including USA) or worse (including China) rating for freedom of press. Proportions of publications originated from countries with problematic or worse situations were 81%, 91%, 61%, 62%, and 63% for political, economic, legislative, sociocultural, and safety/security dimensions, respectively. Results were similar when limited to articles published in 2024 in journals with continuous annual presence in Scopus during 2006-2024. 87.1% of the highly cited papers published in 2024 (with 150 or more Scopus citations by November 23, 2025) have at least one author from a country that is not full democracy and 98.8% of these highly cited papers have at least one author from a country that does not have good freedom of press.

**Conclusions:** Most published science originates from countries struggling or suffering in democracy and/or freedom of press. The deeper causes and implications of this emerging landscape require further study.

## INTRODUCTION

The number of items published every year in the scientific literature is increasing and the landscape of where these publications come from is also changing over time. In a classic analysis published three decades ago in *Science* [1] with the title “The Scientific Wealth of Nations”, data from 1981-1994 demonstrated that 35% of the published scientific papers were published from the USA and these papers received 49% of all citations. Moreover, almost all the literature originated from western democracies. Only India and China were listed among the top 15 most prolific countries, but they published only 2.4% and 0.9% of the literature, respectively, and combined they accounted for just 1% of received citations. Since then, the landscape of published science has changed at a fast pace. China and India currently rank 1^st^ and 3^rd^ worldwide in productivity and impact [2–5].

Social and civil liberties structures have also evolved over time worldwide. Science is a complex, global social phenomenon. Research is published within the existing social, legal, and civil liberty frameworks that prevail across diverse countries. An interesting question is to what extent the scientific literature is currently published by authors in countries that are not full democracies and do not have sufficient freedom of press. There are diverse perspectives about the interaction between democracy and freedom and science [6–10]. Robert Metron, the father of the sociology of science, in his early work believed that science can only flourish in democratic societies [6]. Many authors have since speculated on science and democracy [7–10]. Presence or absence of major civic values may affect what science is prioritized and produced and how it is communicated and disseminated. Science is also increasingly instrumental for policy making [11] and may be invoked or even hijacked to defend decisions with societal and political impact [12]. Societal milieu may shape science-based decision-making and its implementation. Even the reliability of the literature may be affected by the presence or absence of democratic values and freedom of expression. There are mounting concerns about the presence of a replication crisis [13] and decreasing trust in science [14]. Lack of trust is sometimes linked to political interference in scientific matters or the adoption of partisan political endorsements by scientific venues [15,16].

To improve our understanding of the global situation of the scientific literature in relationship to democracy and freedom of press, we aimed to map the origin of published scientific work across the globe in terms of these important civic liberty dimensions.

## METHODS

### Data transparency statement

All data used are made available in the manuscript, publicly accessed sites and its supplements. The presented analyses are descriptive and exploratory, and no protocolized statistical analysis plan was pre-specified. No reporting guideline is pertinent to this type of analysis.

### Democracy and freedom of the press ratings

Democracy ratings in each country worldwide were based on the Democracy Index [17] published annually since 2006 by the Economist Intelligence Unit. The index classifies countries as full democracies (index score >8), flawed democracies (index 6-7.99), hybrid regimes (index 4-5.99), and authoritarian regimes (index <4). The score is obtained by a weighted average of responses to 60 questions in public-opinion surveys that cover electoral process and pluralism, functioning of government, political participation, political culture, and other civil liberties.

Freedom of press ratings are generated annually by Reporters without Borders [18] since 2002, but the currently used scoring approach has been used only since 2022. Overall ratings are classified as good (85-100 points), satisfactory (70-85 points), problematic (55-70 points), difficult (40-55 points) and very serious (0-40 points). Press freedom is defined as “the ability of journalists as individuals and collectives to select, produce, and disseminate news in the public interest independent of political, economic, legal, and social interference and in the absence of threats to their physical and mental safety” [18]. Five separate dimensions are also scored (political, economic, legislative, sociocultural, and safety/security). The index is calculated from the sum of a quantitative tally of recorded abuses and a qualitative analysis from responses of journalists, researchers, academics and human rights defenders to a questionnaire.

It should be acknowledged upfront that the indices used here make normative and methodological assumptions that affect how countries are classified. Furthermore, they may not be interpreted in similar ways in different countries, e.g. China may not perceive itself as an authoritarian regime. Finally, lower index scores should not automatically imply an evaluative judgement about lower scientific quality or legitimacy in the published work of countries that have lower scores.

### Data on scientific publications

The Scopus database [19] was used to obtain (based on data freeze on April 2025) the number of papers published in each country each calendar year. Only items characterized in Scopus as article, review, conference paper, books and book chapter were retained, excluding editorials, notes, letters and other short items. Publication counting was fractionalized by taking the total count of authorships for a paper as the denominator. Thus, a paper authored by author A from China, author B from USA, and author C from USA was counted as 1/3 for China and 2/3 for USA. In the rare cases where an author was affiliated with more than one country, these multiple countries were all counted separately in the denominator. Thus, a paper authored by author A from China and USA, author B from USA and India, and author C from USA was counted as 1/5 for China, 3/5 for USA, and 1/5 for India. The sum of fractional counts was always 1 for each paper. Papers where at least one author had two or more countries were a small minority (5% in 2006, 8% in 2024).

Publications were included regardless of discipline. The National Science Foundation [NSF] has previously used data from Scopus with fractional counts in its Science and Engineering Indicators, trying to limit strictly to science and engineering fields [20]. However, as of May 2025, their released data do not extend beyond 2022 and thus could not be used for analyses pertaining to calendar year 2024. Moreover, inspection of the NSF-used data suggests that they exclude only ∼10% of Scopus-indexed papers anyhow and the excluded papers still represent important scholarly work; boundaries of what disciplines should be included within “science” can be debatable. Therefore, we preferred to be inclusive of all Scopus-covered disciplines.

### Analyses

The analyses are descriptive. Throughout the manuscript, numbers of publications refer to published items count, accounting for fractional assignments of authorships, as detailed above.

First, the countries publishing more than 25,000 publications in 2024 and in 2006 were mapped for their Democracy Index profile in these two calendar years. The countries publishing more than 25,000 publications in 2024 were also mapped for freedom of press.

Second, the count of publications in 2024 and in 2006 was summed for countries in each democracy level and changes in each level were calculated between these two years.

Third, the Pearson correlations between absolute change and relative change in the Democracy Index between 2006 and 2024 and the ratio of publications published in 2024 versus 2006 were estimated.

Fourth, the number of publications published in 2024 was summed for countries in each of the 5 levels of the global Freedom of Press score; and for two major categories (satisfactory of better, problematic or worse) for each of the 5 dimensions of the Freedom of Press score.

Scopus has extended its coverage between 2006 and 2024 with the added indexing of many journals, books and proceedings. One may question whether this expansion has added disproportionately publishing venues that publish a larger share of papers from countries with poor democracy and press freedom ratings. To investigate this possibility, we performed sensitivity analyses, where we considered only publishing venues that have published Scopus-indexed material every year between 2006 and 2024. Besides this “established set”, we also report the data for the publishing venues that first started publishing after 2006 but have been indexed continuously in Scopus every year since they started (“new continuous set”); and the remaining venues (“elliptical set”). The latter set covers both venues that may have been included or discontinued over time in Scopus, as well as venues with intermittent publications, such as conference papers or books.

In all analyses where databases were juxtaposed (Democracy Index versus bibliometrics, Freedom of Press versus bibliometrics), some small countries and territories not represented in both databases were excluded from the calculations. Population data for 2024 are from https://www.populationpyramid.net/population-size-per-country/2024/

Finally, we aimed to appraise the contribution of countries with different ratings regarding democracy and freedom of press towards not only publications, but specifically those publications that receive the highest number of citations. Searches in Scopus were performed on November 23, 2025 to identify for each country the number of papers published in 2024 that had received already at least 150 citations by that date. Of the 4,469,990 items with publication date in 2024, only 1,710 have been so highly cited.

## RESULTS

### Top producing countries

In 2024 there were 27 countries that exceeded 25,000 publications in their annual productivity (Figure 1). Of those, 9 were full democracies, 10 were flawed democracies, 2 were hybrid regimes, and 6 were authoritarian (including the most productive country, China). Conversely in 2006, only 16 countries exceeded 25,000 publications, 9/16 were full democracies (including the leading producer, USA), 5 were flawed democracies, 1 was hybrid regime, and 1 was authoritarian. In 2024, 2/27 top producing countries had good freedom of the press and 8 had satisfactory, while the situation was problematic in 6, difficult in 3, and very serious in 8 (including the leading producer).

**Figure 1.**
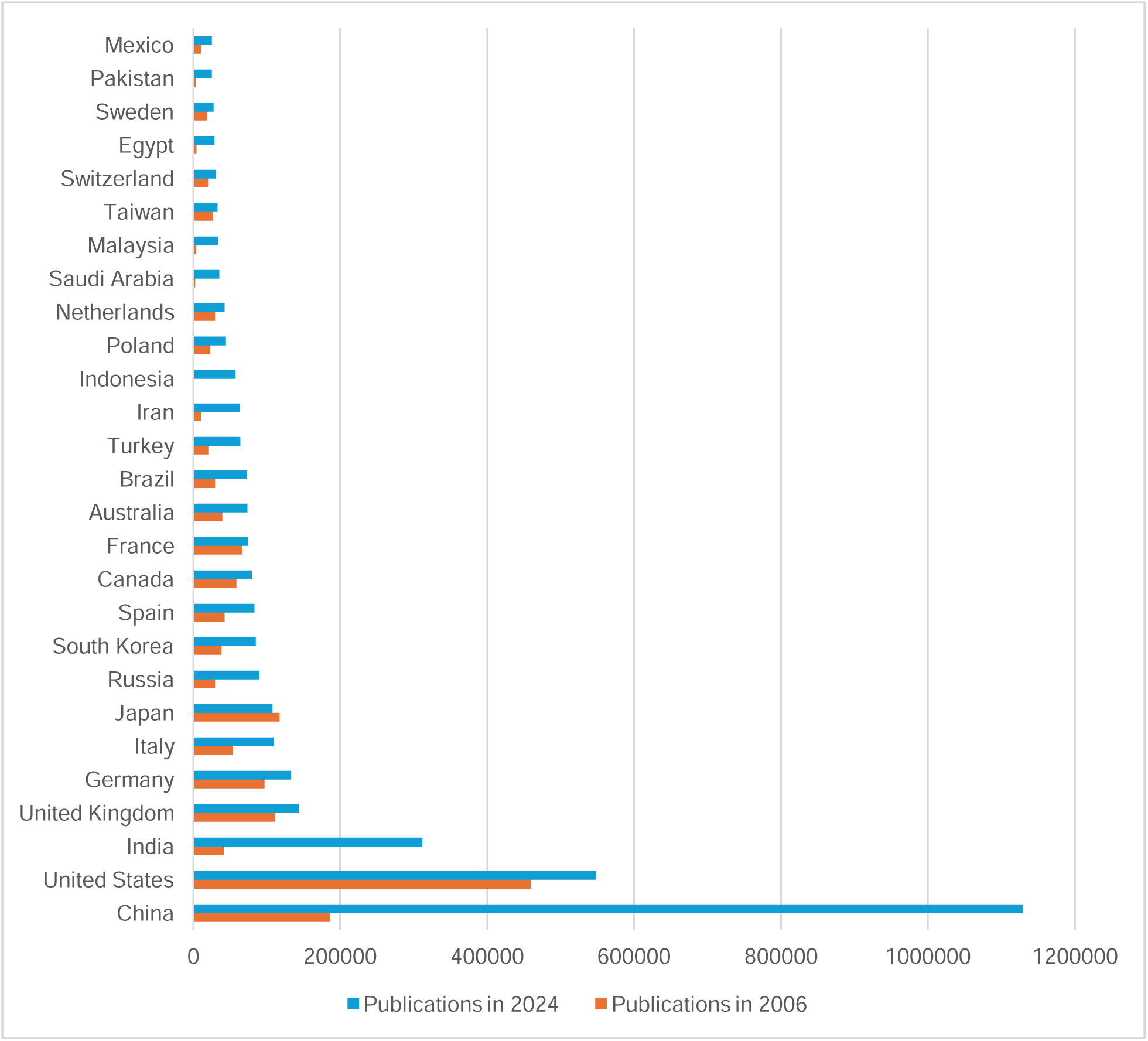
Countries with over 25,000 Scopus-indexed publications in 2024 and comparative data for these countries in 2006.

The top 2 most productive countries had inversed positions between 2006 and 2024. In 2006, USA published 2.5 times more papers than China (459524 versus 186060), while in 2024 China published 2,1 times more papers than the USA (1129320 versus 548584). Both in 2006 and 2024 China was classified as an authoritarian regime and the situation for press freedom in 2024 was characterized as very serious. The USA was classified as full democracy in 2006 but lapsed into flawed democracy ratings since 2016. The USA in 2024 was characterized as problematic in the global index for press freedom; this was true also of the political, economic, and safety dimensions, while it was characterized as satisfactory in the legislative and sociocultural dimensions. In the 2025 ratings released recently by Reporters without Borders [18], also the legislative dimension was scored as problematic.

### Publications and democracy levels in 2006 and 2024

In 2024, 533.0 million people (6,5% of 8,162 total global population) lived in full democracies. As shown in Table 1, the number of countries that were full democracies was almost the same in 2006 and 2024 (26 versus 25), but full democracies accounted for 66% of all publications in 2006 versus only 22% in 2024. Also, the absolute number of publications from full democracies decreased by -21% between 2006 and 2024. Conversely, the proportion of publications accounted by authoritarian regimes almost tripled between 2006 and 2024; this group had the greatest increase in absolute number of publications, a +568% increase.

**Table 1.** Scopus-indexed publications in 2006 and 2024 according to Democracy Index level.

|  | Countries<br>in 2024 | Publications in<br>2024 (%) | Countries<br>in 2006 | Publications in<br>2006 (%) | Change<br>2006-2024 |
| --- | --- | --- | --- | --- | --- |
| **All publications** |  |  |  |  |  |
| Full democracy | 25 | 915102 (22) | 26 | 1157842 (66) | -21% |
| Flawed democracy | 46 | 1552153 (37) | 53 | 301694 (17) | +414% |
| Hybrid regime | 36 | 224526 (5) | 33 | 74155 (4) | +203% |
| Authoritarian regime | 60 | 1494073 (36) | 55 | 223620 (13) | +568% |
| Total | 167 | 4185853 (100) | 167 | 1757310 (100) | +138% |
| **Established set** |  |  |  |  |  |
| Full democracy | 25 | 463487 (23) | 26 | 793929 (66) |  |
| Flawed democracy | 46 | 686884 (35) | 53 | 217333 (17) |  |
| Hybrid regime | 36 | 87423 (4) | 36 | 55473 (4) |  |
| Authoritarian regime | 60 | 751718 (38) | 60 | 156899 (13) |  |
| Total | 167 | 1989511 (100) | 167 | 1223664 (100) |  |
| **New continuous set** |  |  |  |  |  |
| Full democracy | 25 | 289778 (19) | 26 | Not pertinent |  |
| Flawed democracy | 46 | 544863 (36) | 53 | Not pertinent |  |
| Hybrid regime | 36 | 102148 (7) | 33 | Not pertinent |  |
| Authoritarian regime | 60 | 563811 (38) | 55 | Not pertinent |  |
| Total | 167 | 1500600 (100) | 167 | Not pertinent |  |
| **Elliptical set** |  |  |  |  |  |
| Full democracy | 25 | 161837 (23) | 26 | 363913 (68) |  |
| Flawed democracy | 46 | 320405 (46) | 53 | 84361 (16) |  |
| Hybrid regime | 36 | 34956 (5) | 33 | 18682 (4) |  |
| Authoritarian regime | 60 | 178544 (26) | 55 | 66721 (13) |  |
| Total | 167 | 695742 (100) | 167 | 533676 (100) |  |

A number of small countries and territories that have some publications indexed in Scopus but no Democracy Index scoring are not included in the calculations. Scopus data included are available in the Supplement and Democracy Index data are accessible in ref. 17.

67% (1,257,330/1,873,925) of the publications in 2006 and 47% (1,971,218/4,171,333) in 2024 were published by the established set of journals. 35% (1,480,928/4,171,333) of publications in 2024 appeared in the new continuous set of venues. As shown in Table 1, the decline in the share of publications published from full democracies in 2024 versus 2006 and the concomitant increase in the share of publications published from authoritarian regimes was largely similar when the established set, new continuous set, and elliptical set of publishing venues were considered separately. Supplementary Figure 1 shows data for top-producing countries according to publication set.

The decrease in the proportion of papers published between 2006 and 2024 by full democracies is driven almost equally by the much faster acceleration of productivity in countries that were and continue to be not full democracies and by the transition of 2 large, prolific countries (USA and France) from full democracies to flawed democracies. Belgium and Malta also transitioned from full democracies to flawed democracies, while Taiwan, Estonia and Uruguay transitioned from flawed democracies to full democracies. The full democracies of 2006 decreased their share of global literature from 66% to 36% in 2024 (Table 2). Full democracies of 2006 that remained full democracies in 2024 lost two-fifths of their global share (from 35% to 21%), while countries that drifted from full democracies to flawed democracies lost half of their global share (from 31% to 15%) (Table 2).

**Table 2.** Proportion of global Scopus-indexed publications for countries that were full democracies in 2006 and/or 2024.

| Country | Full democracy | Share in 2006, % | Share in 2024, % | Ratio 2024 to 2006 |
| --- | --- | --- | --- | --- |
| United Kingdom | 2006 and 2024 | 6.35 | 3.43 | 0.54 |
| Germany | 2006 and 2024 | 5.53 | 3.17 | 0.57 |
| Japan | 2006 and 2024 | 6.66 | 2.58 | 0.39 |
| Spain | 2006 and 2024 | 2.41 | 1.98 | 0.82 |
| Canada | 2006 and 2024 | 3.32 | 1.90 | 0.57 |
| Australia | 2006 and 2024 | 2.27 | 1.76 | 0.77 |
| Netherlands | 2006 and 2024 | 1.67 | 1.01 | 0.61 |
| Switzerland | 2006 and 2024 | 1.14 | 0.73 | 0.64 |
| Sweden | 2006 and 2024 | 1.05 | 0.66 | 0.63 |
| Portugal | 2006 and 2024 | 0.44 | 0.58 | 1.32 |
| Denmark | 2006 and 2024 | 0.55 | 0.47 | 0.85 |
| Austria | 2006 and 2024 | 0.60 | 0.43 | 0.72 |
| Norway | 2006 and 2024 | 0.46 | 0.42 | 0.92 |
| Greece | 2006 and 2024 | 0.67 | 0.42 | 0.63 |
| Czech Republic | 2006 and 2024 | 0.54 | 0.41 | 0.76 |
| Finland | 2006 and 2024 | 0.65 | 0.38 | 0.59 |
| Ireland | 2006 and 2024 | 0.33 | 0.28 | 0.84 |
| New Zealand | 2006 and 2024 | 0.39 | 0.25 | 0.64 |
| Luxembourg | 2006 and 2024 | 0.02 | 0.04 | 2.22 |
| Iceland | 2006 and 2024 | 0.03 | 0.03 | 1.01 |
| Mauritius | 2006 and 2024 | 0.00 | 0.01 | 2.16 |
| Costa Rica | 2006 and 2024 | 0.02 | 0.02 | 1.64 |
| Taiwan | 2024 only | 1.53 | 0.79 | 0.52 |
| Uruguay | 2024 only | 0.02 | 0.03 | 1.51 |
| Estonia | 2024 only | 0.06 | 0.06 | 1.01 |
| United States | 2006 only | 26.15 | 13.11 | 0.50 |
| France | 2006 only | 3.77 | 1.78 | 0.47 |
| Belgium | 2006 only | 0.86 | 0.54 | 0.62 |
| Malta | 2006 only | 0.01 | 0.02 | 2.61 |

### Correlation between change in publications and change in Democracy Index over time

All countries had higher numbers of publications in 2024 than in 2006 with the exceptions of Venezuela and Japan. The median ratio was 4.38 for publications in 2024 versus those published in 2006. The majority of countries (106/167, 63%) worsened in their Democracy Index between 2006 and 2024 (median change in score, -0.22). For most countries, the change was small. 122 countries did not change democracy overall level, 16 improved by one level, 28 worsened by one level (including the USA), and 1 worsened by two levels. There was no correlation between the ratio of publications in 2024 versus those published in 2006 and the absolute change in the Democracy Index (r=0.02, Figure 2) or the relative change in the Democracy Index (r=0.00) in the same period.

**Figure 2.**
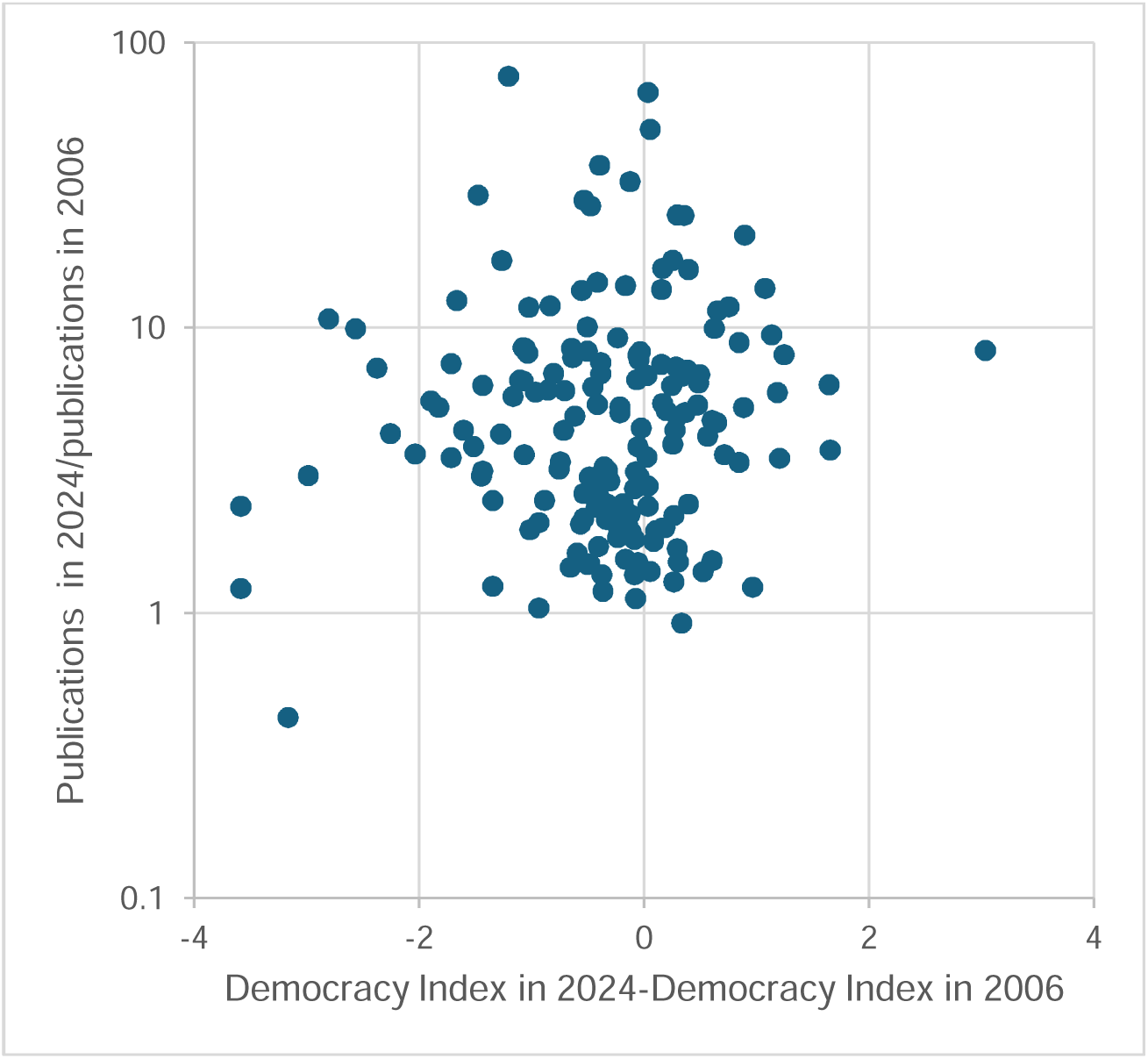
Lack of correlation between ratio of publications in 2024 versus 2006 and absolute change in the Democracy Index.

### Freedom of press and publications in 2024

In 2024, 63.0 million people lived in 8 countries with good freedom of press, and 522.6 million people lived in another 37 countries with satisfactory freedom of press (0.8% and 6.4%, respectively of the 8,162 million total global population). As shown in Table 3, only 4% of publications in 2024 came from countries with good freedom of press and another 18% were from countries with satisfactory freedom of press. More than three-quarters (78%) came from countries with problematic (including USA) or worse (including China) ratings, with most of them coming from countries where the situation for freedom of press was characterized as very serious. Considering separately the 5 dimensions, the proportion of publications in 2024 from countries where the situation was problematic or worse was 81%, 91%, 61%, 62%, and 63% for the political, economic, legislative, sociocultural, and safety/security dimensions, respectively.

**Table 3.** Scopus-indexed publications in 2024 from different countries according to global freedom of press level and its five dimensions.

| Freedom of press level | Countries in 2024 | Publications in 2024, n (%) | Publications in established set, n (%) | Publications in new continuous set, n (%) | Publications in elliptical set, n (%) |
| --- | --- | --- | --- | --- | --- |
| GLOBAL |  |  |  |  |  |
| Good | 8 | 162051 (4) | 82564 (4) | 51646 (3) | 27842 (4) |
| Satisfactory | 37 | 769215 (18) | 385979 (19) | 244719 (16) | 138517 (20) |
| Problematic | 47 | 1116237 (27) | 534691 (27) | 384042 (26) | 197504 (28) |
| Difficult | 49 | 308966 (7) | 104873 (5) | 152766 (10) | 51327 (7) |
| Very serious | 36 | 1830667 (44) | 881763 (44) | 668086 (44) | 280814 (40) |
| Total | 177 | 4187136 (100) | 1989869 (100) | 1501258 (100) | 696008 (100) |
| POLITICAL |  |  |  |  |  |
| At least satisfactory | 27 | 812787 (19) | 415925 (21) | 247106 (16) | 149757 (22) |
| Problematic or worse | 150 | 3374348 (81) | 1573944 (79) | 1254152 (84) | 546251 (78) |
| ECONOMIC |  |  |  |  |  |
| At least satisfactory | 17 | 366325 (9) | 185367 (9) | 115156 (8) | 65802 (9) |
| Problematic or worse | 160 | 3820811 (91) | 1804503 (91) | 1386102 (92) | 630206 (91) |
| LEGISLATIVE |  |  |  |  |  |
| At least satisfactory | 57 | 1649174 (39) | 807391 (41) | 534028 (36) | 307755 (44) |
| Problematic or worse | 120 | 2537962 (61) | 1182478 (59) | 967230 (64) | 388253 (56) |
| SOCIOCULTURAL |  |  |  |  |  |
| At least satisfactory | 59 | 1578334 (38) | 785279 (39) | 496900 (33) | 296155 (43) |
| Problematic or worse | 118 | 2608802 (62) | 1204590 (61) | 1004358 (67) | 399853 (57) |
| **SAFETY/SECURITY** |  |  |  |  |  |
| At least satisfactory | 98 | 1536317 (37) | 716440 (36) | 559987 (37) | 259890 (43) |
| Problematic or worse | 79 | 2650819 (63) | 1273429 (64) | 941271 (63) | 436118 (57) |

A number of small countries and territories that have some publications indexed in Scopus but no Freedom of Press scoring are not included in the calculations. The same applies to OECS, Northern Cyprus, and Kosovo that are rated in Freedom of Press scoring lists but not in Scopus data. Scopus data used are in the Supplement and Freedom of Press data are publicly available in ref. 18.

Results for the established set, the new continuous set and the elliptical set also appear in Table 3 and suggest largely similar patterns across these three sets as in the overall analysis.

### Highly cited papers

Countries that are not full democracies and/or do not have good freedom of press had an overwhelming presence of authors even in the most highly cited papers among those published in 2024 (at least 150 Scopus citations by November 23, 2025) (Table 4). Nevertheless, the proportion of papers that reached such highly cited status among all their papers published tended to be higher in countries with full democracy and good freedom of the press ratings. A total of 3,565,855 papers (77.8% of the total 4,469,990) and 1,489 (87.1% of the total 1,710) highly cited ones had at least one author from a country that was not a full democracy. A total of 4,361,231 papers (97.6% of the total) and 1,690 (98.8% of the total) highly cited ones had at least one author from a country that did not have good freedom of press.

**Table 4.**
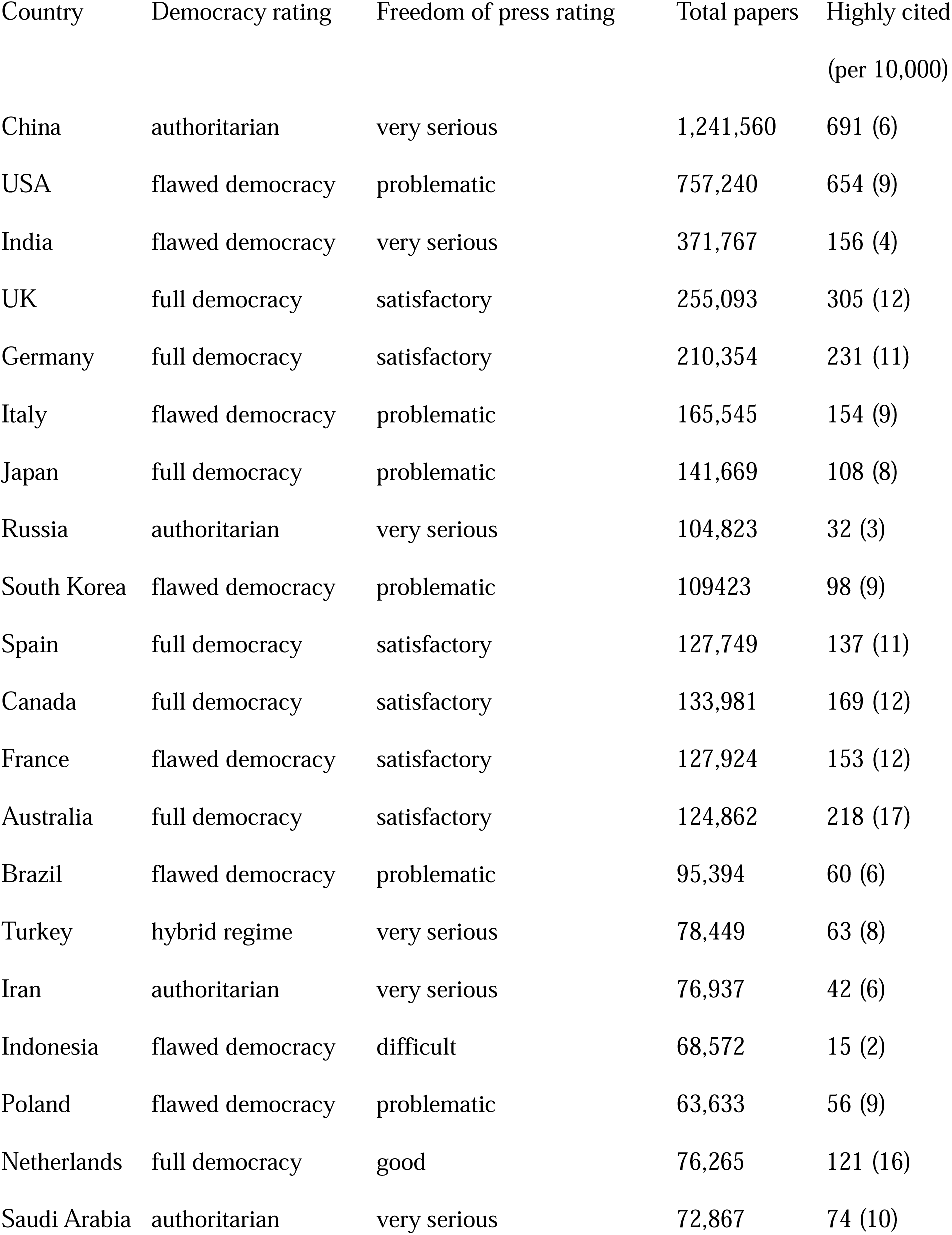
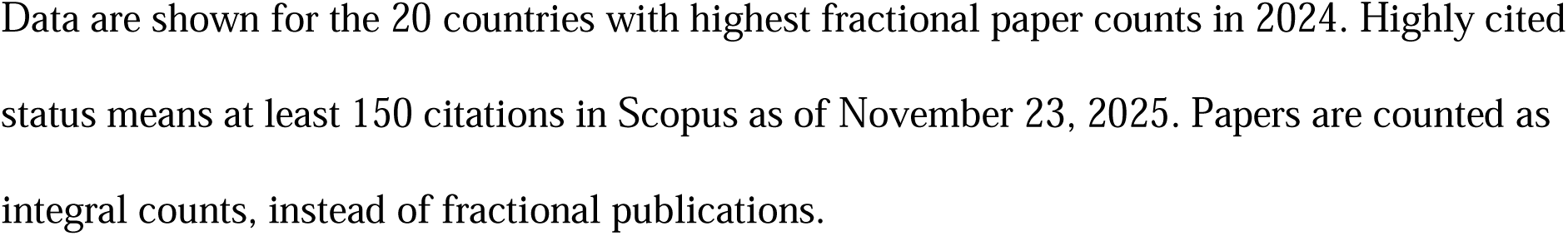
Highly cited papers among those published in 2024.

| Country | Democracy rating | Freedom of press rating | Total papers | Highly cited<br>(per 10,000) |
| --- | --- | --- | --- | --- |
| China | authoritarian | very serious | 1,241,560 | 691 (6) |
| USA | flawed democracy | problematic | 757,240 | 654 (9) |
| India | flawed democracy | very serious | 371,767 | 156 (4) |
| UK | full democracy | satisfactory | 255,093 | 305 (12) |
| Germany | full democracy | satisfactory | 210,354 | 231 (11) |
| Italy | flawed democracy | problematic | 165,545 | 154 (9) |
| Japan | full democracy | problematic | 141,669 | 108 (8) |
| Russia | authoritarian | very serious | 104,823 | 32 (3) |
| South Korea | flawed democracy | problematic | 109423 | 98 (9) |
| Spain | full democracy | satisfactory | 127,749 | 137 (11) |
| Canada | full democracy | satisfactory | 133,981 | 169 (12) |
| France | flawed democracy | satisfactory | 127,924 | 153 (12) |
| Australia | full democracy | satisfactory | 124,862 | 218 (17) |
| Brazil | flawed democracy | problematic | 95,394 | 60 (6) |
| Turkey | hybrid regime | very serious | 78,449 | 63 (8) |
| Iran | authoritarian | very serious | 76,937 | 42 (6) |
| Indonesia | flawed democracy | difficult | 68,572 | 15 (2) |
| Poland | flawed democracy | problematic | 63,633 | 56 (9) |
| Netherlands | full democracy | good | 76,265 | 121 (16) |
| Saudi Arabia | authoritarian | very serious | 72,867 | 74 (10) |

Data are shown for the 20 countries with highest fractional paper counts in 2024. Highly cited status means at least 150 citations in Scopus as of November 23, 2025. Papers are counted as integral counts, instead of fractional publications.

## DISCUSSION

The current analysis shows that more than three quarters of scientific publications in 2024 came from countries without full democracy. This is a complete reversal compared with the late twentieth century [1]. It is also a fundamental change even when compared with 2006 (the first year that Economist Democracy Index ratings started to be issued), when two thirds of publications were published from countries with full democracy. There was no correlation between changes in democracy ratings and changes in scientific productivity between 2006 and 2024 across countries. Moreover, in 2024 more than three quarters of scientific papers were published from countries with problematic or worse status for freedom of press.

Our analyses also show that among papers published in 2024, the proportion of those that have reached highly cited status by November 2025 tends to be lower for countries that are not full democracies and/or have no good freedom of press. However, given the sheer volume of publications from such countries, they still occupy the top rankings even in the production of highly cited papers in terms of absolute numbers. Moreover, 87.1% of the highly cited papers have at least one author from a country that is not full democracy and 98.8% of highly cited papers have at least one author from a country that does not have good freedom of press.

The major underlying changes in these patterns are the rapid, major growth in the scientific productivity of China [2–4], a country classified as an authoritarian regime and with very serious situation regarding freedom of press; the decline in democracy rating and the problematic performance in freedom of press of the USA; and the extreme growth in Scopus-indexed papers in several other countries that are not full democracies. In 2014 China published more than twice the number of Scopus-indexed papers than the USA and the gap keeps widening. Moreover, since 2021 it published more papers in the top 1% of citations or in the select group of journals captured by the Nature Index [2]. The USA retains superiority only at the very top of the most extremely cited papers (top 100 published in a year) [21]. India (a country classified as flawed democracy and with very serious situation for freedom of press) has increased its annual Scopus-indexed publications by almost 8-fold between 2006 and 2024. If it continues at this pace, it may also exceed the USA by 2029 [5]. Several countries with authoritarian regimes and very serious situation for freedom of press, such as Russia, Turkey, Iran, Saudi Arabia, and Pakistan have had 3-16-fold the Scopus-indexed scientific publications in 2024 than in 2006. Indonesia, a country characterized as flawed democracy with difficult situation for freedom of press has accelerated 67-fold its annual Scopus-indexed publications during this time.

We explored also the possibility of indexing bias as Scopus has become more inclusive over time. When analyses were limited to journals that have published Scopus-indexed material every year between 2006-2024, we saw a similar pattern of emerging dominance of countries with struggling democracy and freedom press ratings. Concurrently, we observed that a substantial share of the Scopus-indexed literature is currently published in venues that started publishing in the last 2 decades and the dominance of countries with struggling democracy and freedom press rankings is largely similar also in these venues. A major growth of the volume of publications in this time period has been due to mega-journals that publish massively thousands of papers every year, in particular with special issues (22,23). There is ongoing debate about these journals (23,24). Regardless, they seem to cater disproportionately to authors from some countries, in particular less developed countries (e.g. in the Middle East) and Eastern Europe (25-27).

The implications of these emerging patterns can be debated. On the one hand, democracy and freedom of press do not appear to be necessary to publish many scientific papers. Full democracies and countries with good or satisfactory freedom of press accounted in 2024 for a 3-5-fold larger share of the published literature than their respective share of the global population. However, they represent a very small portion of the global population. Their higher per capita number of Scopus-indexed publications may be related more to their typically higher financial resources rather than their civil liberties.

Changes in the geography of science may reflect broader transformations in state capacity, industrial policy, and internationalisation rather than a decline of democratic values. Many countries with major increases in Scopus-indexed publications have purposefully supported this growth. Many non-democratic states have deliberately invested in research as part of national modernisation strategies or to strengthen their global influence. On the one hand, some authoritarian regimes have even used questionable aggressive incentives to propel scientific productivity [28,29]. On a positive note, however, higher attention to and investment in science may be positive both for science and for society at large. E.g. China has largely caught up with the USA in overall R&D investment [30] and has surpassed the USA in number of patents and number of new businesses formed [31,32]. Chinese scientists who trained in the USA and returned to China may also have been instrumental in this positive evolution [33,34]. Countries with increasing openness to foreign collaboration may benefit the most in terms of productivity and impact growth [35]. Moreover, as a nation-state, China has enacted policies that may allow rapid scientific growth, e.g. protection of intellectual property, encouraging mobility and skills development, and government procurement of science and technology, including military purchases [36]. Previous work has suggested that less democratic countries in the international system are reaping gains in scientific performance through increases in external complexity (globalization and international collaboration) [37].

Conversely, one may worry that less democratic and more authoritarian regimes may shape more aggressively what scientific topics are preferred or even allowed to be pursued. They may also have higher levels of interference in science, especially for matters that may have political, economic, or societal repercussions. Political interference in science is a hot topic even for countries with strong past democratic traditions and views can become polarized, fueling further extremes of partisanship and parallel discussions and concerns about decline in democracy. Moreover, aggressive incentives for publications may result in overproduction of pointless papers and even fraud, e.g. paper mills and citation cartels [38–39]. Some questionable and unethical publishing behaviors are indeed becoming more frequent among authors from these countries with massive increases in productivity [40], resulting in high retraction rates [41]. This may be the tip of the iceberg of a large problem with the evolving reliability of the scientific literature. Conversely, this may be a biased assessment, if papers from such countries are considered more suspect because of cultural bias and easier to retract.

There can also be debate of whether and how freedom of press might affect the scientific literature. Most influences, if any, may be indirect, or freedom of the press may be a surrogate marker of other civil and cultural values that are instrumental for conducting and disseminating science. Journalism, and in particular science journalism, is crucial in disseminating science and its inferences to the broader public. For science that becomes politicized, lack of press freedom can have grave consequences. Moreover, science may be more easily hijacked for various political or related agendas without the press being able to provide critical, unbiased commentary. Biased, partisan journalism may even be instrumental for facilitating and whitewashing the hijacking of science.

The contemporary research ecosystem is very complex, and democracy or lack thereof may interfere in positive or negative ways with many factors that shape that ecosystem. These include national research priorities and the tolerance for topics outside of or even apparently contradicting national priorities and preferred narratives; funding structures and the balance between public, industry, and other funding sources; organizational models; recruitment practices (including corruption levels) and their control or lack thereof by different government, private, institutional or local stakeholders; research training and research evaluation systems and any expectations that they may nudge for or impose; and many others. Democratic and other civil liberties permeate the research ecosystem and how it functions, and shape what it is expected to produce.

Moreover, the dynamics of the journal publication system should be considered. The number of Scopus-indexed journals and annual publications has increased substantially between 2006 and 2024. The growth of total publication volume has seen a 7% average year on year (136% over the 19 years), while for the fixed set of journals that published consistently between 2006-2024, the average growth rate is 3.6% year on year (69% over the entire period). In terms of the geographic location of the publishers, these continue to be mostly in western countries, at least for those journals that are indexed by Scopus. E.g. even the new flagship of Chinese science, the Innovation, aimed to be a competitor to Science and Nature, is published by Cell Press. However, this affords no guarantee of quality of the peer review. Peer review is overfatigued and it is difficult to secure qualified peer reviewers [42]. It is likely that many poor papers get accepted without substantive peer review even outside of predatory journals that perform no review at all [43]. Even positive changes such as the wide availability of free datasets may backfire, if these are used for massive production of low-quality and flawed papers by authors who lack the necessary expertise [44]. Furthermore, the COVID-19 pandemic may have created additional strain to the system in the quest for massive, fast publications [45,46] and this may have left persistent dis-regulation even long after the end of the pandemic.

## LIMITATIONS

Some limitations of the current analyses should be discussed. First, the democracy and freedom of press indices used here are complex constructs that may not fully capture with perfect accuracy the intended entities. Different people may have different perspectives. Some may even challenge whether these indices may have explicit bias favoring western countries with liberal or neoliberal economies. However, these indices are widely used. For similar (but not identical) constructs, additional indices are also available. Classification of countries would be modestly different with different indices. For example, the Freedom in the World rating [48] has less granularity in classification, using only three categories: free, partly free, not free. Therefore, it lists (as of 2025) 83 of 194 countries and territories as free [49], but freedom carries large variability within this category. Another index related to societal civil values, the Corruption Perceptions Index [50] has a granular classification in increments of 10 points at a 0-100 scale (100 suggesting zero corruption). Based on this index, only 8 countries worldwide have protection from corruption score >80 and they publish only a tiny proportion of the scientific literature.

Second, no effort was made to break down the bibliographic production to diverse scientific fields. Analyses of finer granularity may be pursued in the future by those interested in specific disciplines. Research in some subdisciplines may be strongly disincentivized or even banned in specific countries and regimes. However, focusing on between-fields difference may miss the big picture of the radical overall change described here.

Third, one should be cautious in making too bold causal statements about the link between a political regime and research integrity and misconduct. Given the complex ecosystems of both politics and science, highly specific causality is very difficult to establish.

Fourth, we focused on publication numbers and provided some data also on highly cited papers, but research quality and rigor may not be properly captured with such indices. For methodological quality and rigor, it is very difficult to generalize across studies of very different designs and topics.

## CONCLUSIONS

Acknowledging these caveats, there has been a clear, fundamental change in the scientific literature landscape, and the global scientific output has become increasingly concentrated over time in countries with lower democracy and press freedom scores. The causes, implications, and potential need for any interventions for this potentially disquieting phenomenon require attention and further study.

## DECLARATIONS

### Ethics approval and consent to participate

Not applicable

### Consent for publication

Not applicable

### Availability of data and materials

All data are in the manuscript and supplement

## Funding

None

### Authors’ contributions

JPAI had the idea, collected data, analyzed data, and wrote the paper. JB contributed to the concept discussion, collected data, and edited the paper. Both authors approved the final version.

## Acknowledgements

Not applicable

## Supplementary Data

Number of publications in Scopus in 2006 and 2024 (fractional count)

| Country | 2006 | 2024 | 2006<br>Established | 2024<br>Established | 2024 New<br>continuous | 2006<br>Elliptical | 2024<br>Elliptical |
| --- | --- | --- | --- | --- | --- | --- | --- |
| China | 186,060.3 | 1,129,320.2 | 132,447.5 | 609,824.7 | 387,612.4 | 53,612.8 | 131,883.1 |
| United States | 459,524.2 | 548,583.9 | 309,732.5 | 282,289.0 | 150,321.9 | 149,791.7 | 115,973.0 |
| India | 41,162.3 | 311,561.4 | 30,206.1 | 110,446.0 | 103,035.5 | 10,956.2 | 98,079.8 |
| United Kingdom | 111,586.5 | 143,768.1 | 76,654.4 | 71,892.0 | 39,780.6 | 34,932.0 | 32,095.5 |
| Germany | 97,194.1 | 132,663.1 | 65,098.5 | 66,769.5 | 40,459.1 | 32,095.7 | 25,434.6 |
| Italy | 54,058.7 | 109,431.2 | 38,245.3 | 49,833.0 | 41,619.4 | 15,813.5 | 17,978.8 |
| Japan | 117,026.8 | 108,010.8 | 78,623.0 | 57,567.8 | 33,078.5 | 38,403.7 | 17,364.6 |
| Russia | 29,549.3 | 89,783.2 | 23,884.4 | 33,142.6 | 44,023.3 | 5,664.9 | 12,617.3 |
| South Korea | 38,399.2 | 85,238.2 | 28,505.4 | 37,849.9 | 37,596.8 | 9,893.8 | 9,791.5 |
| Spain | 42,342.2 | 82,937.5 | 31,969.9 | 37,462.7 | 35,468.8 | 10,372.3 | 10,006.0 |
| Canada | 58,421.1 | 79,641.5 | 40,351.3 | 42,190.4 | 22,867.7 | 18,069.8 | 14,583.4 |
| France | 66,332.3 | 74,688.9 | 45,518.5 | 41,007.1 | 20,714.8 | 20,813.9 | 12,967.0 |
| Australia | 39,843.0 | 73,488.6 | 27,819.9 | 40,455.6 | 21,043.6 | 12,023.1 | 11,989.4 |
| Brazil | 29,468.8 | 73,303.7 | 22,593.1 | 38,332.8 | 27,370.8 | 6,875.7 | 7,600.1 |
| Turkey | 20,412.1 | 64,299.9 | 15,631.5 | 33,631.9 | 22,607.3 | 4,780.6 | 8,060.7 |
| Iran | 10,688.7 | 63,764.0 | 6,924.2 | 30,168.6 | 28,786.9 | 3,764.5 | 4,808.6 |
| Indonesia | 858.2 | 57,597.3 | 505.9 | 13,169.4 | 35,497.9 | 352.3 | 8,929.9 |
| Poland | 22,911.8 | 44,575.1 | 16,450.2 | 18,434.1 | 20,690.4 | 6,461.5 | 5,450.7 |
| Netherlands | 29,319.6 | 42,483.3 | 21,060.8 | 23,119.9 | 12,191.2 | 8,258.7 | 7,172.3 |
| Saudi Arabia | 2,191.5 | 35,514.7 | 1,455.4 | 14,083.8 | 17,731.9 | 736.1 | 3,699.0 |
| Malaysia | 3,555.5 | 33,685.1 | 1,939.4 | 10,578.0 | 17,390.7 | 1,616.1 | 5,716.4 |
| Taiwan | 26,876.6 | 33,175.7 | 18,533.2 | 17,166.7 | 10,894.7 | 8,343.4 | 5,114.3 |
| Switzerland | 20,063.6 | 30,357.9 | 14,248.2 | 15,662.4 | 9,410.9 | 5,815.4 | 5,284.7 |
| Egypt | 4,354.3 | 28,607.1 | 3,009.2 | 13,233.4 | 12,029.6 | 1,345.2 | 3,344.1 |
| Sweden | 18,528.9 | 27,714.3 | 13,900.5 | 14,671.1 | 8,514.0 | 4,628.3 | 4,529.3 |
| Pakistan | 2,979.6 | 25,437.0 | 2,109.6 | 12,172.9 | 10,148.5 | 870.0 | 3,115.6 |
| Mexico | 10,098.7 | 25,087.1 | 7,075.3 | 10,534.2 | 10,879.1 | 3,023.4 | 3,673.9 |
| Portugal | 7,814.8 | 24,483.0 | 5,388.5 | 9,509.2 | 11,212.0 | 2,426.3 | 3,761.8 |
| South Africa | 7,001.2 | 23,749.2 | 5,068.4 | 9,085.3 | 9,860.2 | 1,932.8 | 4,803.7 |
| Belgium | 15,103.9 | 22,419.2 | 10,546.1 | 12,079.6 | 6,748.2 | 4,557.8 | 3,591.4 |
| Thailand | 4,709.6 | 22,336.3 | 3,218.0 | 7,051.6 | 12,228.4 | 1,491.5 | 3,056.3 |
| Iraq | 279.8 | 21,392.9 | 151.1 | 6,506.0 | 12,304.9 | 128.7 | 2,582.0 |
| Hong Kong | 10,145.3 | 21,083.5 | 7,143.3 | 11,239.5 | 6,071.5 | 3,002.0 | 3,772.6 |
| Ukraine | 5,426.4 | 19,638.1 | 3,675.2 | 4,659.3 | 12,259.1 | 1,751.2 | 2,719.7 |
| Denmark | 9,600.1 | 19,508.4 | 7,237.7 | 10,908.4 | 5,540.7 | 2,362.4 | 3,059.3 |
| Austria | 10,516.6 | 18,024.3 | 7,197.8 | 8,840.7 | 5,807.5 | 3,318.8 | 3,376.1 |
| Norway | 8,079.6 | 17,784.0 | 5,938.5 | 9,379.9 | 5,299.1 | 2,141.1 | 3,105.1 |
| Greece | 11,783.6 | 17,730.5 | 8,434.3 | 6,655.1 | 8,005.9 | 3,349.3 | 3,069.5 |
| Israel | 12,640.4 | 17,679.3 | 9,519.9 | 9,699.2 | 5,189.2 | 3,120.4 | 2,790.9 |
| Czech Republic | 9,487.4 | 17,234.9 | 6,742.3 | 8,352.2 | 6,494.5 | 2,745.1 | 2,388.2 |
| Vietnam | 495.1 | 16,166.4 | 346.0 | 5,492.1 | 8,132.6 | 149.2 | 2,541.7 |
| Finland | 11,450.6 | 16,044.7 | 7,637.6 | 8,261.4 | 4,519.5 | 3,813.0 | 3,263.8 |
| Singapore | 9,289.4 | 15,636.8 | 6,220.8 | 7,296.2 | 4,691.9 | 3,068.6 | 3,648.7 |
| Romania | 4,135.1 | 14,880.2 | 2,413.6 | 4,070.4 | 8,270.7 | 1,721.5 | 2,539.1 |
| Morocco | 1,070.8 | 14,763.3 | 722.4 | 4,441.8 | 7,985.0 | 348.5 | 2,336.5 |
| Nigeria | 2,911.1 | 13,575.8 | 1,365.5 | 3,739.8 | 6,954.5 | 1,545.6 | 2,881.5 |
| Chile | 3,292.4 | 12,639.9 | 2,522.7 | 5,337.8 | 5,894.9 | 769.8 | 1,407.1 |
| Colombia | 1,598.4 | 12,396.5 | 1,160.9 | 4,051.0 | 6,683.5 | 437.5 | 1,662.0 |
| Hungary | 5,922.0 | 11,620.8 | 4,232.8 | 4,505.7 | 5,200.9 | 1,689.2 | 1,914.1 |
| Ireland | 5,791.2 | 11,550.7 | 3,966.2 | 5,763.1 | 3,394.7 | 1,825.0 | 2,393.0 |
| Argentina | 6,003.8 | 11,539.2 | 4,506.8 | 5,711.8 | 4,291.6 | 1,497.0 | 1,535.9 |
| United Arab Emirates | 971.5 | 11,183.5 | 526.4 | 3,482.6 | 4,518.5 | 445.1 | 3,182.5 |
| Bangladesh | 887.1 | 11,084.9 | 491.0 | 2,814.5 | 5,267.2 | 396.1 | 3,003.1 |
| New Zealand | 6,799.7 | 10,392.1 | 5,026.1 | 5,609.3 | 2,789.1 | 1,773.6 | 1,993.7 |
| Algeria | 1,433.9 | 10,209.6 | 731.6 | 3,928.7 | 4,250.4 | 702.3 | 2,030.5 |
| Ethiopia | 311.2 | 9,116.4 | 244.7 | 3,075.9 | 5,509.8 | 66.5 | 530.7 |
| Jordan | 1,061.8 | 8,369.2 | 742.3 | 2,457.2 | 4,802.5 | 319.5 | 1,109.5 |
| Peru | 576.1 | 8,339.4 | 476.2 | 1,650.8 | 5,517.7 | 99.8 | 1,170.9 |
| Tunisia | 2,078.0 | 7,764.0 | 1,363.4 | 3,044.7 | 3,520.9 | 714.6 | 1,198.4 |
| Serbia | 2,358.1 | 7,686.2 | 1,582.1 | 2,842.5 | 3,604.0 | 776.0 | 1,239.7 |
| Philippines | 532.3 | 7,270.8 | 379.1 | 1,541.0 | 3,213.4 | 153.2 | 2,516.3 |
| Croatia | 3,245.5 | 6,989.4 | 2,180.2 | 2,291.1 | 3,457.9 | 1,065.3 | 1,240.5 |
| Kazakhstan | 242.3 | 6,811.2 | 190.6 | 1,510.7 | 4,427.8 | 51.7 | 872.7 |
| Slovakia | 2,787.0 | 6,754.9 | 2,141.5 | 2,354.7 | 3,302.0 | 645.4 | 1,098.1 |
| Bulgaria | 2,021.6 | 6,471.3 | 1,336.2 | 1,738.2 | 2,939.0 | 685.4 | 1,794.1 |
| Uzbekistan | 334.1 | 5,779.2 | 275.5 | 1,773.1 | 3,377.4 | 58.6 | 628.7 |
| Ecuador | 151.9 | 5,661.4 | 105.9 | 1,231.9 | 3,703.6 | 45.9 | 725.8 |
| Slovenia | 2,709.1 | 5,253.4 | 1,996.9 | 2,102.1 | 2,232.2 | 712.2 | 919.1 |
| Ghana | 238.8 | 5,059.7 | 167.8 | 1,700.4 | 2,673.0 | 71.0 | 686.2 |
| Lithuania | 1,968.9 | 3,918.5 | 1,458.2 | 1,247.5 | 2,195.2 | 510.8 | 475.8 |
| Macao | 206.9 | 3,852.5 | 118.0 | 1,945.8 | 1,301.0 | 88.9 | 605.7 |
| Qatar | 214.2 | 3,440.0 | 112.8 | 1,241.1 | 1,475.8 | 101.4 | 723.0 |
| Lebanon | 761.3 | 3,248.5 | 537.7 | 1,430.8 | 1,429.7 | 223.6 | 388.0 |
| Sri Lanka | 444.0 | 3,073.2 | 249.4 | 851.2 | 1,166.8 | 194.6 | 1,055.2 |
| Oman | 405.6 | 2,962.8 | 296.7 | 808.2 | 1,385.3 | 108.9 | 769.4 |
| Cyprus | 553.3 | 2,793.6 | 334.6 | 1,210.9 | 1,035.6 | 218.7 | 547.0 |
| Kenya | 606.1 | 2,705.0 | 459.6 | 883.8 | 1,375.0 | 146.6 | 446.2 |
| Azerbaijan | 314.1 | 2,608.0 | 225.4 | 728.3 | 1,538.0 | 88.7 | 341.7 |
| Estonia | 1,028.7 | 2,482.6 | 694.4 | 950.8 | 974.5 | 334.3 | 557.3 |
| Nepal | 389.4 | 2,314.5 | 273.1 | 880.5 | 1,153.7 | 116.3 | 280.3 |
| Tanzania | 326.9 | 2,229.2 | 239.3 | 766.6 | 1,192.8 | 87.6 | 269.9 |
| Kuwait | 760.0 | 2,209.7 | 504.5 | 755.9 | 1,044.9 | 255.5 | 409.0 |
| Cameroon | 361.4 | 2,180.7 | 268.1 | 940.1 | 1,060.7 | 93.3 | 179.9 |
| Latvia | 431.2 | 2,112.7 | 258.9 | 485.5 | 1,200.5 | 172.3 | 426.7 |
| Uganda | 230.5 | 1,959.0 | 189.7 | 782.3 | 893.3 | 40.7 | 283.4 |
| Luxembourg | 331.3 | 1,753.7 | 178.7 | 724.2 | 515.0 | 152.7 | 514.4 |
| Belarus | 1,257.3 | 1,563.0 | 999.4 | 497.4 | 839.5 | 257.9 | 226.1 |
| Cuba | 1,445.8 | 1,507.7 | 948.4 | 611.7 | 760.7 | 497.4 | 135.3 |
| Uruguay | 394.2 | 1,417.8 | 303.4 | 658.7 | 504.9 | 90.8 | 254.3 |
| Palestinian Territory | 142.5 | 1,417.8 | 105.7 | 451.9 | 798.1 | 36.8 | 167.9 |
| Bosnia and Herzegovina | 320.1 | 1,402.0 | 195.0 | 414.3 | 732.3 | 125.1 | 255.5 |
| Armenia | 396.6 | 1,386.1 | 302.1 | 555.8 | 600.8 | 94.5 | 229.5 |
| Bahrain | 194.6 | 1,269.7 | 139.8 | 259.3 | 546.9 | 54.8 | 463.4 |
| Georgia | 507.7 | 1,186.2 | 398.7 | 395.8 | 531.3 | 109.0 | 259.2 |
| Iceland | 444.6 | 1,071.3 | 343.6 | 520.5 | 344.4 | 101.0 | 206.5 |
| Costa Rica | 263.8 | 1,032.7 | 209.9 | 322.4 | 529.6 | 53.9 | 180.6 |
| Zimbabwe | 196.9 | 995.5 | 136.3 | 235.2 | 407.3 | 60.7 | 353.0 |
| Syria | 116.7 | 953.1 | 79.8 | 258.0 | 588.2 | 36.9 | 106.9 |
| Albania | 34.5 | 862.4 | 22.2 | 124.3 | 590.4 | 12.4 | 147.7 |
| Sudan | 122.7 | 772.8 | 84.7 | 248.9 | 383.4 | 38.0 | 140.6 |
| Malta | 124.2 | 772.1 | 59.5 | 237.0 | 312.8 | 64.6 | 222.2 |
| Kyrgyzstan | 55.4 | 750.8 | 38.3 | 120.5 | 544.5 | 17.1 | 85.8 |
| North Macedonia | 246.7 | 743.4 | 127.6 | 168.8 | 396.2 | 119.0 | 178.4 |
| Yemen | 59.1 | 699.3 | 45.7 | 222.8 | 357.8 | 13.4 | 118.6 |
| Senegal | 163.3 | 682.1 | 77.9 | 224.6 | 360.5 | 85.4 | 96.9 |
| Brunei | 47.9 | 658.9 | 32.5 | 167.2 | 295.7 | 15.4 | 196.0 |
| Venezuela | 1,511.5 | 651.2 | 890.9 | 195.4 | 330.3 | 620.6 | 125.5 |
| Burkina Faso | 112.7 | 649.4 | 80.1 | 229.1 | 342.7 | 32.6 | 77.6 |
| Jamaica | 390.4 | 633.9 | 210.9 | 217.2 | 184.3 | 179.5 | 232.3 |
| Puerto Rico | 691.4 | 631.0 | 486.3 | 269.7 | 264.3 | 205.1 | 96.9 |
| Rwanda | 23.1 | 619.7 | 15.9 | 213.6 | 336.5 | 7.2 | 69.5 |
| Libya | 110.3 | 592.8 | 56.4 | 135.2 | 249.6 | 53.9 | 208.0 |
| Malawi | 110.3 | 581.2 | 86.8 | 200.4 | 258.7 | 23.5 | 122.1 |
| Benin | 76.4 | 573.6 | 44.6 | 198.5 | 316.5 | 31.8 | 58.7 |
| Botswana | 239.5 | 567.9 | 162.9 | 201.4 | 241.2 | 76.6 | 125.3 |
| Zambia | 82.3 | 528.5 | 67.5 | 178.6 | 264.0 | 14.8 | 85.8 |
| Moldova | 173.5 | 480.4 | 136.1 | 164.7 | 238.6 | 37.4 | 77.1 |
| Ivory Coast | 137.9 | 466.1 | 73.4 | 158.3 | 261.2 | 64.5 | 46.5 |
| Democratic Republic of the Congo | 36.7 | 439.3 | 19.7 | 181.5 | 225.3 | 17.0 | 32.5 |
| Mongolia | 60.0 | 395.9 | 38.6 | 151.1 | 184.6 | 21.4 | 60.2 |
| Panama | 37.2 | 375.3 | 28.1 | 83.2 | 164.4 | 9.1 | 127.7 |
| Namibia | 46.7 | 374.8 | 38.3 | 114.5 | 133.2 | 8.4 | 127.1 |
| Paraguay | 39.3 | 363.3 | 20.1 | 99.9 | 210.2 | 19.2 | 53.2 |
| Somalia | 2.0 | 352.5 | 2.0 | 77.7 | 254.0 | - | 20.8 |
| Montenegro | 64.7 | 352.3 | 23.5 | 141.5 | 131.6 | 41.2 | 79.1 |
| Mauritius | 68.1 | 350.4 | 44.8 | 73.4 | 137.7 | 23.3 | 139.3 |
| Bolivia | 89.8 | 315.6 | 70.4 | 95.5 | 152.2 | 19.4 | 67.8 |
| Afghanistan | 26.7 | 287.5 | 13.9 | 68.3 | 181.7 | 12.8 | 37.5 |
| Cambodia | 52.9 | 278.9 | 38.9 | 89.1 | 140.5 | 14.0 | 49.3 |
| Mozambique | 48.8 | 271.2 | 41.9 | 112.4 | 128.2 | 6.9 | 30.6 |
| Honduras | 14.8 | 254.7 | 10.4 | 36.7 | 160.1 | 4.4 | 57.8 |
| Fiji | 107.6 | 251.3 | 64.3 | 90.4 | 123.5 | 43.3 | 37.4 |
| Togo | 30.4 | 245.4 | 14.3 | 85.6 | 148.1 | 16.2 | 11.7 |
| Myanmar | 34.3 | 238.0 | 18.8 | 61.1 | 59.4 | 15.5 | 117.5 |
| Reunion | 82.3 | 226.5 | 56.9 | 123.7 | 80.5 | 25.4 | 22.3 |
| North Korea | 4.5 | 222.2 | 3.6 | 139.4 | 71.1 | 0.8 | 11.7 |
| Madagascar | 60.2 | 180.9 | 46.7 | 74.8 | 89.0 | 13.5 | 17.1 |
| Guatemala | 44.9 | 172.7 | 30.3 | 60.0 | 80.9 | 14.7 | 31.7 |
| Tajikistan | 33.5 | 164.8 | 28.9 | 51.1 | 93.0 | 4.7 | 20.7 |
| Dominican Republic | 23.9 | 164.4 | 17.0 | 58.5 | 86.3 | 7.0 | 19.6 |
| Mali | 65.9 | 156.5 | 39.1 | 59.6 | 71.7 | 26.8 | 25.1 |
| Bhutan | 18.4 | 153.7 | 13.2 | 44.7 | 77.3 | 5.2 | 31.6 |
| Gabon | 53.5 | 140.7 | 36.3 | 58.2 | 72.1 | 17.2 | 10.4 |
| Sierra Leone | 11.6 | 138.3 | 9.5 | 52.9 | 58.4 | 2.2 | 27.0 |
| Niger | 30.7 | 130.4 | 18.2 | 35.6 | 76.8 | 12.5 | 18.0 |
| Swaziland | 39.3 | 124.1 | 27.5 | 45.2 | 53.4 | 11.8 | 25.5 |
| Papua New Guinea | 60.0 | 123.2 | 34.8 | 48.3 | 47.9 | 25.2 | 27.1 |
| Republic of the Congo | 48.0 | 121.5 | 29.1 | 45.9 | 55.1 | 18.9 | 20.5 |
| El Salvador | 25.7 | 112.6 | 18.8 | 23.5 | 42.5 | 6.9 | 46.5 |
| Guyana | 11.4 | 94.6 | 5.9 | 21.8 | 40.5 | 5.5 | 32.3 |
| Angola | 14.9 | 94.1 | 6.3 | 32.0 | 48.5 | 8.5 | 13.6 |
| Laos | 39.7 | 93.0 | 32.1 | 35.3 | 47.0 | 7.6 | 10.8 |
| Lesotho | 16.5 | 89.0 | 13.9 | 22.6 | 35.0 | 2.6 | 31.4 |
| Gambia | 49.2 | 87.9 | 46.1 | 33.8 | 42.7 | 3.1 | 11.4 |
| Liberia | 3.5 | 86.7 | 2.0 | 31.8 | 37.9 | 1.5 | 17.1 |
| New Caledonia | 64.3 | 86.1 | 61.2 | 51.3 | 21.4 | 3.1 | 13.4 |
| French Guiana | 27.0 | 85.1 | 19.8 | 38.9 | 38.9 | 7.1 | 7.3 |
| Trinidad and Tobago | 29.6 | 81.0 | 14.5 | 19.6 | 25.0 | 15.2 | 36.4 |
| Mauritania | 9.1 | 80.6 | 7.5 | 23.6 | 44.8 | 1.6 | 12.2 |
| Greenland | 20.1 | 75.4 | 18.1 | 40.8 | 18.1 | 2.1 | 16.5 |
| Liechtenstein | 40.6 | 73.2 | 25.5 | 31.4 | 24.1 | 15.1 | 17.7 |
| Grenada | 17.2 | 72.9 | 16.8 | 34.4 | 32.6 | 0.3 | 5.9 |
| Guinea | 20.4 | 72.0 | 13.6 | 26.2 | 36.7 | 6.8 | 9.1 |
| Turkmenistan | 5.1 | 71.8 | 2.8 | 12.6 | 44.1 | 2.3 | 15.1 |
| Chad | 10.6 | 66.7 | 10.0 | 21.1 | 36.1 | 0.6 | 9.5 |
| Monaco | 24.3 | 66.3 | 18.6 | 36.4 | 20.9 | 5.7 | 9.0 |
| Faroe Islands | 5.7 | 65.7 | 4.7 | 38.2 | 19.9 | 1.1 | 7.6 |
| Maldives | 5.0 | 63.8 | 3.0 | 30.3 | 24.1 | 2.0 | 9.3 |
| Nicaragua | 48.3 | 58.9 | 34.7 | 23.0 | 33.5 | 13.6 | 2.4 |
| Burundi | 7.6 | 55.0 | 5.5 | 17.1 | 28.9 | 2.1 | 8.9 |
| Eritrea | 24.9 | 53.2 | 21.6 | 17.9 | 24.7 | 3.3 | 10.6 |
| Haiti | 16.7 | 50.6 | 13.7 | 26.6 | 18.3 | 3.0 | 5.7 |
| Vatican | 10.3 | 50.2 | 6.8 | 11.8 | 27.8 | 3.5 | 10.6 |
| Guam | 34.3 | 47.4 | 17.9 | 24.9 | 19.8 | 16.4 | 2.7 |
| French Polynesia | 17.4 | 45.3 | 14.7 | 28.1 | 13.4 | 2.7 | 3.9 |
| Saint Kitts and Nevis | 5.7 | 44.0 | 4.7 | 19.1 | 22.3 | 1.0 | 2.6 |
| South Sudan | 1.0 | 43.7 | 1.0 | 10.8 | 28.5 | - | 4.5 |
| East Timor | 4.0 | 39.7 | 2.6 | 13.4 | 23.6 | 1.4 | 2.7 |
| Cape Verde | 5.0 | 37.1 | 3.6 | 13.6 | 20.4 | 1.3 | 3.1 |
| Barbados | 28.4 | 35.8 | 14.8 | 12.8 | 14.8 | 13.6 | 8.2 |
| Central African Republic | 14.5 | 34.2 | 8.0 | 15.0 | 13.9 | 6.5 | 5.3 |
| Bahamas | 5.7 | 31.5 | 4.8 | 14.8 | 9.2 | 0.9 | 7.6 |
| Martinique | 24.3 | 30.1 | 14.6 | 17.3 | 11.3 | 9.7 | 1.5 |
| San Marino | 9.9 | 28.1 | 1.9 | 10.8 | 12.3 | 8.0 | 5.0 |
| Djibouti | 4.0 | 27.4 | 2.4 | 8.5 | 13.7 | 1.7 | 5.2 |
| Seychelles | 10.0 | 26.6 | 8.5 | 13.8 | 9.9 | 1.5 | 2.9 |
| Guadeloupe | 22.8 | 24.7 | 14.3 | 11.2 | 11.4 | 8.5 | 2.1 |
| U.S. Virgin Islands | 14.6 | 24.3 | 9.7 | 15.2 | 6.7 | 4.9 | 2.4 |
| Guinea-Bissau | 8.5 | 23.8 | 8.5 | 12.2 | 11.6 | - | - |
| Samoa | 9.6 | 23.7 | 4.6 | 10.3 | 8.0 | 5.0 | 5.4 |
| Suriname | 5.2 | 22.8 | 4.9 | 7.9 | 8.4 | 0.3 | 6.4 |
| Belize | 4.8 | 22.7 | 4.8 | 10.7 | 4.6 | - | 7.4 |
| Vanuatu | 5.8 | 19.8 | 4.3 | 7.7 | 7.3 | 1.5 | 4.7 |
| Curacao | - | 19.0 | 6.2 | 11.0 | 1.8 |  |  |
| Solomon Islands | 7.6 | 18.7 | 5.1 | 9.5 | 7.7 | 2.5 | 1.6 |
| Antigua and Barbuda | 3.0 | 16.5 | 0.1 | 5.6 | 6.7 | 2.9 | 4.2 |
| Gibraltar | 3.4 | 16.4 | 1.3 | 7.4 | 8.1 | 2.1 | 0.9 |
| Sao Tome and Principe | 1.0 | 14.2 | 1.0 | 7.4 | 5.9 | - | 0.9 |
| Cayman Islands | 10.1 | 13.8 | 5.8 | 5.9 | 7.6 | 4.3 | 0.2 |
| Bermuda | 17.0 | 12.6 | 15.1 | 7.2 | 4.8 | 1.8 | 0.7 |
| Andorra | 7.2 | 12.1 | 5.2 | 2.9 | 7.0 | 2.0 | 2.2 |
| Comoros | 1.4 | 11.6 | 0.2 | 6.2 | 3.4 | 1.2 | 2.0 |
| Aruba | - | 10.6 | 6.6 | 4.0 | - |  |  |
| Micronesia | 3.1 | 10.5 | 3.1 | 1.5 | 2.6 | - | 6.3 |
| Jersey | 8.7 | 9.4 | 7.8 | 3.9 | 2.4 | 0.9 | 3.1 |
| Anguilla | 0.2 | 7.9 | 0.2 | 2.1 | 5.7 | - | - |
| Palau | 6.8 | 7.2 | 2.8 | 5.1 | 1.5 | 4.0 | 0.6 |
| Dominica | 2.1 | 7.1 | 1.1 | 2.4 | 4.1 | 1.0 | 0.7 |
| American Samoa | 4.0 | 6.3 | 4.0 | 1.8 | 2.8 | - | 1.8 |
| Falkland Islands | 11.2 | 6.3 | 10.2 | 3.1 | 2.1 | 1.0 | 1.0 |
| Equatorial Guinea | 4.0 | 6.2 | 3.2 | 3.5 | 1.6 | 0.8 | 1.1 |
| Saint Vincent and the Grenadines | 0.7 | 5.5 | 0.7 | 2.6 | 0.2 | - | 2.7 |
| Tonga | 4.7 | 5.1 | 1.7 | 3.3 | 1.8 | 3.0 | - |
| United States Minor Outlying Islands | - | 5.0 | - | - | 5.0 |  |  |
| Netherlands Antilles | 12.3 | 4.3 | 11.6 | 2.8 | 1.6 | 0.8 | - |
| Montserrat | 3.2 | 4.3 | 2.3 | 3.9 | 0.4 | 1.0 | - |
| Saint Lucia | 4.0 | 4.0 | 2.6 | 2.8 | 1.0 | 1.3 | 0.2 |
| Cook Islands | 0.7 | 3.2 | 0.7 | 0.4 | 2.7 | - | 0.1 |
| British Virgin Islands | 3.1 | 3.2 | 2.1 | 0.1 | 1.3 | 1.0 | 1.8 |
| Mayotte | 3.8 | 3.2 | 1.5 | 1.1 | 2.1 | 2.3 | 0.1 |
| Kiribati | 0.9 | 2.9 | 0.4 | 0.5 | 1.1 | 0.5 | 1.2 |
| Norfolk Island | - | 2.6 | 0.3 | 0.3 | 2.0 |  |  |
| Saint Helena | 0.1 | 2.6 | 0.1 | 1.6 | 0.1 | - | 0.9 |
| British Indian Ocean Territory | - | 2.4 | 2.1 | 0.1 | 0.2 |  |  |
| Niue | 2.0 | 1.7 | - | 1.2 | 0.5 | 2.0 | - |
| South Georgia and the South Sandwich Islands | - | 1.7 | - | 0.6 | 1.1 |  |  |
| Tuvalu | 0.8 | 1.6 | 0.8 | 0.6 | 1.0 | - | - |
| Turks and Caicos Islands | 1.8 | 1.5 | 1.3 | 0.6 | 0.9 | 0.5 | - |
| Northern Mariana Islands | 0.8 | 1.5 | 0.8 | 1.0 | 0.5 | - | - |
| Marshall Islands | 4.1 | 1.3 | 3.6 | 0.2 | 1.1 | 0.5 | - |
| Serbia and Montenegro | 17.4 | 1.2 | 16.4 | 1.0 | 0.2 | 1.0 | - |
| Bouvet Island | - | 1.0 | - | 1.0 | - |  |  |
| Sint Maarten | - | 0.9 | 0.3 | 0.6 | - |  |  |
| Svalbard and Jan Mayen | - | 0.9 | 0.9 | - | - |  |  |
| Antarctica | 1.5 | 0.9 | 0.5 | 0.0 | 0.5 | 1.0 | 0.3 |
| Pitcairn | - | 0.7 | - | 0.7 | - |  |  |
| Saint Martin | - | 0.5 | 0.1 | 0.0 | 0.4 |  |  |
| Tokelau | - | 0.5 | 0.5 | - | - |  |  |
| Saint Pierre and Miquelon | - | 0.5 | - | 0.5 | - |  |  |
| Nauru | 0.3 | 0.0 | 0.3 | 0.0 | - | - | - |
| Cocos Islands | 0.8 | - | - |  | 0.8 |  |  |
| Christmas Island | 0.5 | - | 0.5 |  | - |  |  |

**Supplementary Figure 1.**
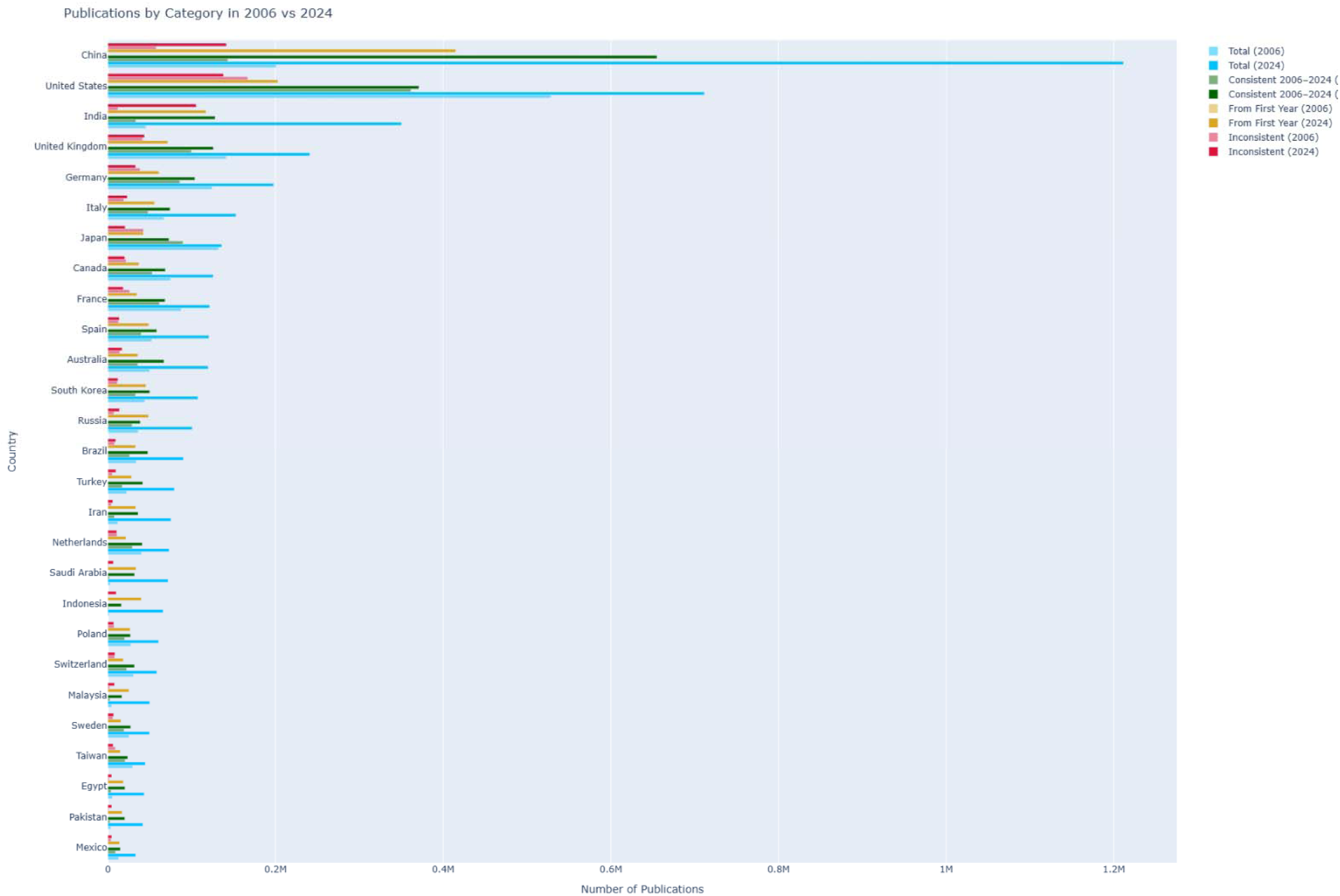
Top-producing countries with fractional count of publications in 2006 and 2024 broken down according to type of venue

